# rsofun v5.0: A model-data integration framework for simulating ecosystem processes

**DOI:** 10.1101/2023.11.24.568574

**Authors:** Josefa Arán Paredes, Koen Hufkens, Mayeul Marcadella, Fabian Bernhard, Benjamin D. Stocker

**Affiliations:** Institute of Geography, University of Bern, Hallerstrasse 12, 3012 Bern, Switzerland; Oeschger Center for Climate Change Research (OCCR), University of Bern, Hochschulstrasse 6, 3012 Bern, Switzerland

## Abstract

Mechanistic vegetation models serve to estimate terrestrial carbon fluxes and climate impacts on ecosystems across diverse biotic and abiotic conditions. Systematically informing them with data is key for enhancing their predictive accuracy and estimating uncertainty. Here we present the Simulating Optimal FUNctioning {rsofun} R package, providing a computationally efficient and parallelizable implementation of the P-model for site-scale simulations of ecosystem photosynthesis, complemented with functionalities for Bayesian model-data integration and estimation of parameters and uncertainty. We describe a use case to demonstrate the package functionalities for modelling ecosystem gross CO_2_ uptake at one flux measurement site, including model sensitivity analysis, Bayesian parameter calibration, and prediction uncertainty estimation. {rsofun} lowers the bar of entry to ecosystem modelling and model-data integration and serves as an open-access resource for model development and dissemination.

## 1 Introduction

The modelling of land ecosystem processes and structure, water, and carbon fluxes relies on both mechanistic and statistical approaches (Dietze et al., 2018; Hartig et al., 2012; Van Oijen et al., 2005). Mechanistic models are formulated as mathematical descriptions of functional relationships between the abiotic environment and ecosystem states, rates, and dynamics. These descriptions reflect available theory and general empirical patterns and provide a means for “translating” hypotheses about governing principles and causal relationships into testable predictions (Marquet et al., 2014), and for upscaling model-based estimates in geographical space and to novel environmental conditions. However, mechanistic models rely also on statistical (empirical) descriptions of processes at varying levels of abstraction.

Mechanistic models have model parameters that are either specified directly or fitted to data. A great advantage of mechanistic models is that they explicitly link known physical constants with process representations (e.g., molecular mass of CO_2_ for diffusion and assimilation, or the gravitational constant and viscosity of water for its transport and transpiration). Other parameters may be specified based on independent measurements under controlled conditions (e.g., the activation energy of Arrhenius-type metabolic rates), or represent measurable plant functional traits, taken as constant over time and within plant functional types (PFTs - the basic unit in mechanistic vegetation models). Both types of parameters have traditionally been specified in models directly (‘direct parameterization’, (Hartig et al., 2012)). Yet other model parameters may not be directly observable and describe processes that are not explicitly resolved but can be described at a higher level of abstraction. Such parameters are often fitted to data such that the agreement between one (or several) related model predictions and observations is optimised. Parameter estimation for mechanistic vegetation models typically employs generic optimization algorithms or Bayesian statistical approaches and is often used for specifying diverse types of parameters (except for universal physical constants). Bayesian methods have the advantage that they enable a systematic assessment of the correlation structure among multiple fitted parameters, provide a means for considering uncertainty in observations and available prior information, and generate probabilistic parameter estimations and model predictions (Dietze et al., 2018; Hartig et al., 2012; Van Oijen et al., 2005).

As the number of parameters increases in state-of-the-art mechanistic vegetation models, taking into account multiple PFTs and ecosystem components (e.g. soil, microbes, hydrology), larger amounts of data and computing resources are required to fully explore the parameter space (Hartig et al., 2012). This poses a limitation for systematic model-data integration and Bayesian parameter estimation. Eco-evolutionary optimality (EEO) principles have been proposed for reducing model complexity and for a robust grounding of models in governing principles (Franklin et al., 2020; Harrison et al., 2021). They enable parameter-sparse representations, limit the distinction of separate PFTs, and may enable better model generalisations to novel environmental regimes. EEO principles make predictions of plant functional traits that would otherwise have to be prescribed – typically as temporally fixed model parameters. However, although parameter-sparse, EEO-based vegetation models are not devoid of model parameters. Ideally, remaining parameters represent known physical constants or quantities that can be measured independently. But remaining parameters in optimality models typically also represent quantities that are not directly measurable – e.g., the marginal cost of water in (Cowan and Farquhar, 1977), or the unit cost ratio in (Prentice et al., 2014). These are considered to be more universally valid (e.g., without distinctions between PFTs), but still must be fitted to data.

The P-model (Prentice et al., 2014; Stocker et al., 2020; Wang et al., 2017) is an example of an EEO-guided model for terrestrial photosynthesis and its acclimation. It avoids the requirement for prescribing PFT-specific parameters of photosynthesis and stomatal regulation but instead predicts them from universal EEO principles for the full range of environmental conditions across the Earth’s (C_3_ photosynthesis-dominated) biomes. However, not directly observable parameters of the underlying EEO theory and of additional empirical parameterizations employed in the P-model (Stocker et al., 2020) remain and must be specified or fitted to data (Tab. 2).

Here, we provide a solution for this challenge, acknowledging that the data is an integral part of the modelling process (Dietze et al., 2013) - even in theory-based models of ecosystem processes. We show how the most important parameters contributing to uncertainty in the P-model can be estimated using observations of ecosystem-level photosynthetic CO_2_ uptake (gross primary productivity, GPP). We provide a brief description of the theory embodied in the P-model and introduce the P-model implementation in the Simulating Optimal FUNctioning {rsofun} version v5.0 modelling framework, made available as an R package. A more comprehensive P-model description can be found in (Stocker et al., 2020). We demonstrate the functionalities of {rsofun} through a case study for simulating GPP at a single flux measurement site. The case study includes sensitivity analysis, Bayesian model calibration, and inference – the prediction of GPP with an estimation of its uncertainty. This paper presents the calibration to GPP observations only, but the package allows calibration to multiple targets simultaneously, including fluxes and leaf traits.

## 2 P-model description

The P-model predicts the acclimation of leaf-level photosynthesis to a (slowly varying) environment based on EEO principles. It thereby yields a parameter-sparse representation of ecosystem-level quantities, generalising across (C_3_ photosynthesis-dominated) vegetation types and biomes. The P-model combines established theory for C_3_ photosynthesis following the Farquhar-von Caemmerer-Berry (FvCB, (Farquhar et al., 1980)) model with the Least-Cost hypothesis for the optimal balancing of water loss and carbon gain (Prentice et al., 2014), and the coordination hypothesis (Wang et al., 2017), which states that the light and Rubisco-limited assimilation rates (as described by the FvCB model) are equal for representative daytime environmental conditions. Based on these theoretical foundations, gross primary productivity (GPP) can be modelled as the product of absorbed photosynthetically active radiation, specified by the locally measured photosynthetically active radiation (PAR) and remotely sensed fAPAR, and the theory-based prediction of the ecosystem-level light use efficiency (LUE) (Stocker et al., 2020; Wang et al., 2017). LUE acclimates to preceding environmental conditions with a characteristic, empirically determined (calibrated), time scale *τ*. Two latent (not directly observable) parameters govern the optimality-guided water-carbon trade-off: the unit cost ratio *β* (governing the balancing of maintaining the carboxylation capacity versus the transpiration stream) and *c*^*^ (the marginal cost of maintaining the electron transport rate). These have previously been calibrated separately to data and specified as fixed model parameters in the P-model (Stocker et al., 2020; Wang et al., 2017). The theory for predicting acclimated LUE then requires only the atmospheric environment to be specified (meteorological variables and CO_2_). A set of corollary predictions, physically and physiologically consistent with the simulated LUE, follows. These include the acclimated base rates of photosynthetic capacities in the FvCB model (*V*_cmax25_ and *J*_max25_), the acclimated average ratio of leaf internal-to-ambient CO_2_ concentration (*c*_i_:*c*_a_), acclimated average daytime stomatal conductance (*g*_s_), and the acclimated base rate of leaf dark respiration (*R*_d25_). Physical constants and additional parameters that determine the instantaneous temperature dependence of *V*_cmax_, *J*_max_, *R*_d_, and parameters in the FvCB model are prescribed and held fixed in the P-model (Tab. A2 in (Stocker et al., 2020)).

For simulating GPP, the P-model is conceived as a single-big-leaf model (Fig. 1). While describing leaf-level quantities at relatively high mechanistic detail, the link between the leaf and the canopy-scale is not explicitly resolved. Instead, an empirical approach for leaf-to-canopy scaling of CO_2_ uptake is employed by treating the quantum yield parameter *φ*_0_ to be representative for the canopy-scale and allowing it to be calibrated to ecosystem-level CO_2_ flux data. The implementation of the P-model in the {rsofun} package (version v5.0) further includes an empirical parameterization of the temperature dependency of the quantum yield *φ*_0_, generalising the approach taken in (Stocker et al., 2020)), and an empirical soil moisture stress function, generalising the approach taken in (Stocker et al., 2020)) (Appendix A). Acclimating quantities are derived by employing the P-model theory to gradually varying environmental conditions where variations are damped and lagged by a characteristic (calibrated) time scale *τ*. The continuous treatment of the acclimation time scale is different from (Stocker et al., 2020), where monthly mean values of environmental variables were considered for the acclimation. The temperature dependency of *φ*_0_ can be turned off by setting *a* (see Tab. 2 and Appendix A) to 0 (‘ORG’ model setup in (Stocker et al., 2020)). The soil moisture stress function can be turned off by setting *β*_0_ (see Tab. 2 and Appendix A) to 1 (‘BRC’ model setup in (Stocker et al., 2020)). Taken together, the P-model approach is guided by EEO and thus yields predictions for quantities that otherwise must be prescribed and fitted to data. Yet, a small set of model parameters, related to empirical parametrizations and latent quantities remain and must be calibrated to data. See Tab. 2 for a list of all (calibratable) model parameters of the P-model implementation in the {rsofun} package.

**Figure 1:**
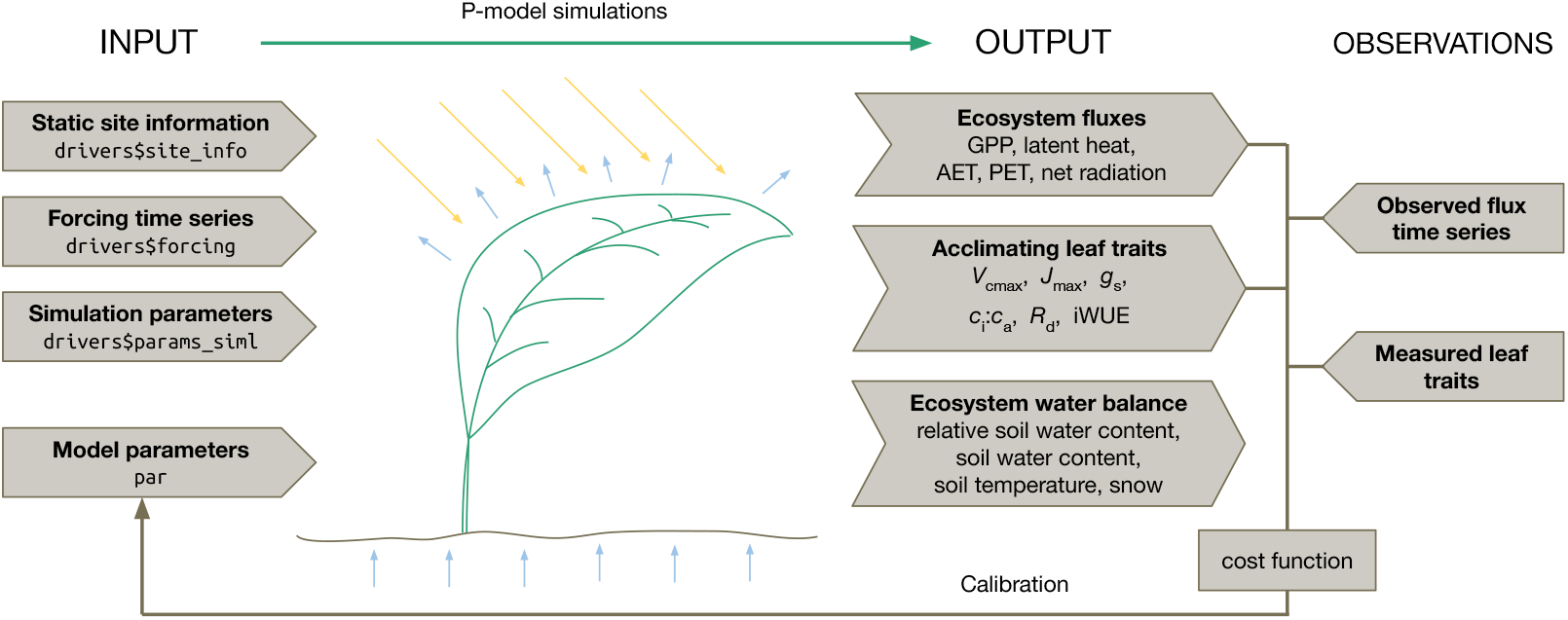
P-model inputs, outputs, and target observations for the calibration. The model takes as inputs static site information (longitude, latitude, elevation and root zone water storage capacity), time series of meteorological forcings (listed in Tab. 1), simulation parameters (spinup years, recycle period length, vegetation type (evergreen, deciduous, grass (C3/C4) with/without N-fixation) and, common for all sites, model parameters (listed in Tab. 2). The simulation returns a time series of several ecosystem fluxes, acclimating leaf traits and ecosystem water balance quantities. By comparing these outputs to field measurements and flux data in a Bayesian calibration routine, model parameters can be estimated.

## 3 The {rsofun} model framework

{rsofun} implements the P-model (Stocker et al., 2020) and provides off-the-shelf methods for Bayesian (probabilistic) parameter and prediction uncertainty estimation. {rsofun} is distributed as an R package on R’s central and public package repository. {rsofun} also implements the BiomeE vegetation demography model (Weng et al., 2019, 2015). The latter is not further described here and is implemented at an experimental stage in {rsofun} version v5.0. The P-model implementation in {rsofun} is designed for time series simulations by accounting for temporal dependencies in the acclimation to a continuously varying environment (Tab. 1). Function wrappers in R make the simulation workflow user-friendly and all functions and input forcing data structures are comprehensively documented (https://geco-bern.github.io/rsofun/).

**Table 1:**
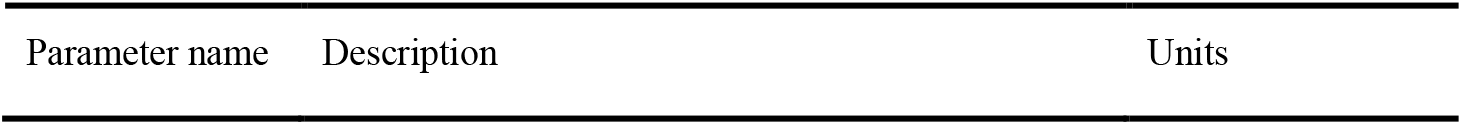
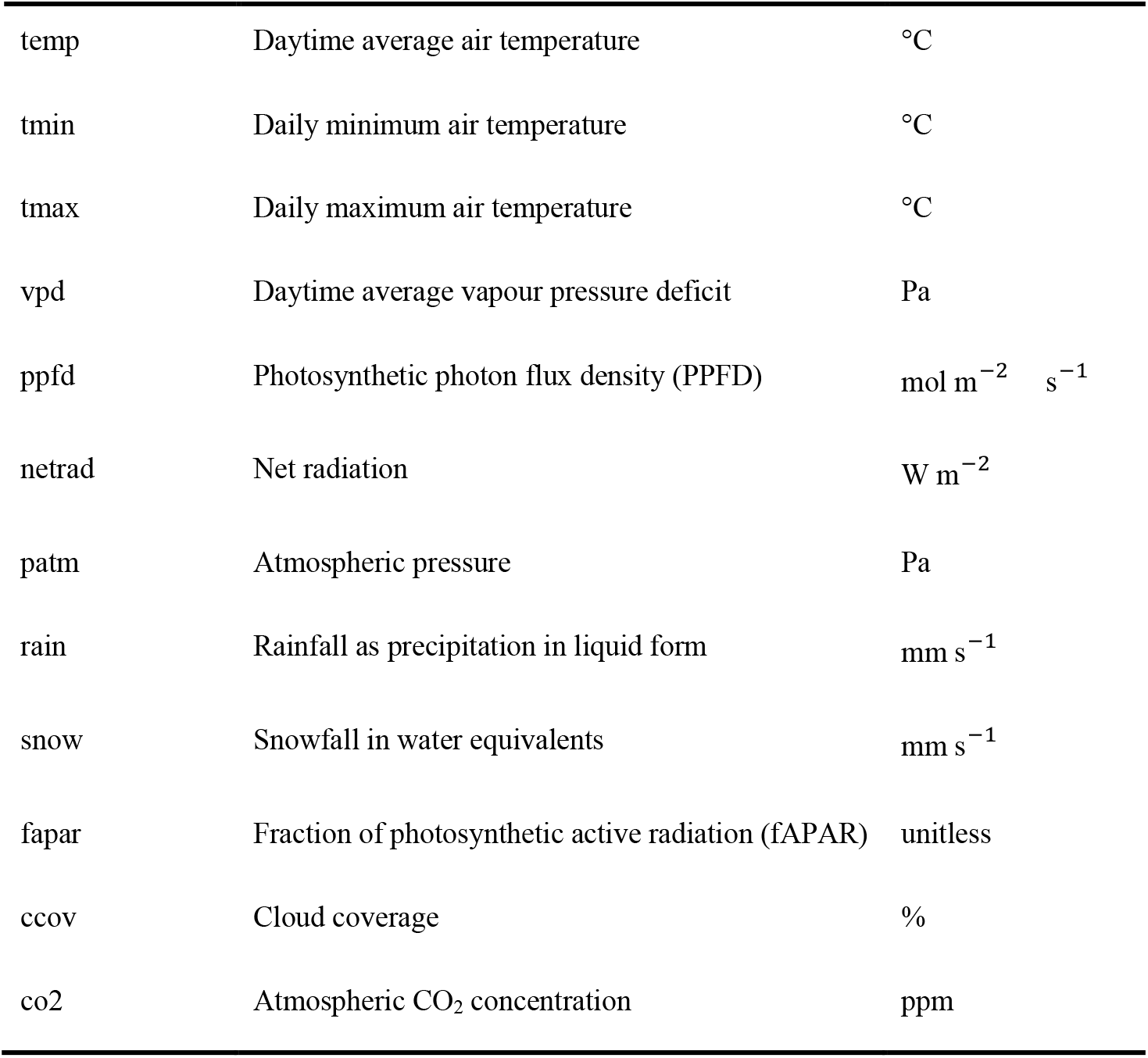
Forcing time series. Daily time series of the following meteorological variables are required for simulation. If a spin-up period is specified, the corresponding years are recycled for the spin-up.

In {rsofun}, model parameters can be calibrated using a calibration function ‘calib_sofun()’, providing two modes of calibration, one based on generalised simulated annealing ({GenSA} R package) for global optimization (Xiang et al., 2013) and one based on Markov chain Monte Carlo (MCMC) implemented by the {BayesianTools} R package, giving access to a wide variety of Bayesian methods (Hartig et al., 2023). The former being fast, while the latter provides more informed parameter optimization statistics (Clark, 2004; Dietze et al., 2013). This gives the option for both exploratory and more in-depth analysis of estimated parameters. A set of standard cost functions are provided for the calibration, facilitating the exploration of various metrics or target variables and the specification of calibrated model parameters. Furthermore, the vignettes accompanying the package (https://geco-bern.github.io/rsofun/articles/) explain how to customise the calibration cost functions and interpret the calibration results.

## 4 Case study

We use {rsofun} to model GPP as estimated from ecosystem flux measurements taken at one site using the eddy-covariance technique. The site, selected here for demonstration purposes, is Puéchabon (Rambal et al., 2004), an evergreen Mediterranean forest, dominated by *Quercus ilex*, growing on relatively shallow soil on karstic bedrock, and located in southern France. The climate is governed by a distinct seasonality in solar radiation and temperature, peaking in summer, and a dry period during summer months with recurrent ecosystem water limitation in late summer (Rambal et al., 2004). Data used here cover years 2007-2012. Both the model target data (GPP) and model forcing data (see Fig. 1) were measured at the site and obtained from the FLUXNET2015 dataset (Pastorello et al., 2021). Data for years 2013 and 2014 were available from FLUXNET2015 but removed here due to unexpectedly low GPP measurements (*pers. comm. Jean-Marc Limousin*).

### 4.1 Sensitivity analysis

The P-model implementation in {rsofun} has a total of nine model parameters that are available for calibration (Tab. 2). Due to the relatively high computational cost of simultaneously calibrating all nine model parameters, we start by performing a sensitivity analysis to determine the most influential parameters on the model fit and exclude the least influential ones for the subsequent calibration step. Here, we apply the Morris method for global sensitivity analysis (Morris, 1991) from the {sensitivity} R package (Looss et al., 2023) and compute the sensitivity metrics *μ*\*, indicating the magnitude of the overall influence of a given parameter on the prediction, and *σ*, a measure of the heterogeneity of a parameter’s influence on the prediction across the parameter space. We assume an additive and normally distributed model error term for the GPP prediction by the P-model (Trotsiuk et al., 2020) and express the fit to observed data via the Gaussian log-likelihood. The sensitivity analysis result (Fig. 2) indicates that the quantum yield intercept parameter *φ*_0_ is the most influential parameter, followed by the unit cost of electron transport *c\** and the optimal temperature for the quantum yield *b*. The remaining parameters have relatively little influence on the evaluation of daily GPP predictions from the single site considered here and *b*_0_, the ratio of dark respiration to the temperature-normalised maximum carboxylation rate, has no influence on GPP predictions, which follows from the model structure. Additional analyses (not shown here) indicated that the convergence of the parameter calibration (shown in the next section) is undermined when calibrating *β* and *c*^*^ simultaneously with other model parameters. Therefore, and based on the sensitivity analysis (Fig. 2), we chose to hold *c*^*^, β, τ, and *b*_0_ constant and thereby excluded them from the Bayesian calibration procedure described in the next section.

**Table 2:** Calibratable parameters in the {rsofun} P-model implementation. The maximum a posteriori (MAP) estimates from the Bayesian model calibration (Sec. 4.2) are listed in column ‘Values’. The bounds of uniform prior distributions are given in parentheses. Parameters that were held fixed for the calibration are marked with an asterisk (*). Fixed site information was: longitude = 3.6°E, latitude = 43.7°N, elevation = 270 m a.s.l., and total root zone water storage capacity = 432 mm.

| Symbol | Parameter name | Description | Units | Values |
| --- | --- | --- | --- | --- |
| $\varphi_0$ | kphio | Quantum yield at optimal temperature | mol mol<sup>-1</sup> | 0.041<br>(0.02, 0.15) |
| $a$ | kphio_par_a | Shape parameter for the temperature dependence of the quantum yield | °C<sup>-2</sup> | -0.0023<br>(-0.004, -0.001) |
| $b$ | kphio_par_b | Optimal temperature for the quantum yield | °C | 15.3<br>(10, 30) |
| $\theta^*$ | soilm_thetastar | Threshold plant-available soil water content in the soil moisture stress function | mm | 158<br>(4.3, 432) |
| $\beta_0$ | soilm_betao | Stress factor at low soil moisture, intercept for the soil moisture stress function | unitless | 0.0001<br>(0, 1) |
| $\beta$ | beta_unitcostratio | Unit cost ratio of carboxylation (maintenance of $V_{\text{cmax}}$) to transpiration | unitless | 146* |
| $b_0$ | rd_to_vcmax | Ratio of temperature-normalised dark respiration ($R_{\text{d25}}$) to the temperature-normalised maximum carboxylation rate $V_{\text{cmax25}}$ (eq. C8 in Stocker et al. 2020) | unitless | 0.014* |
| $\tau$ | tau_acclim | Acclimation time scale of photosynthesis | days | 20* |
| $c^*$ | kc_jmax | Unit cost of electron transport (maintenance of $J_{\text{max}}$) | unitless | 0.41* |

**Figure 2:**
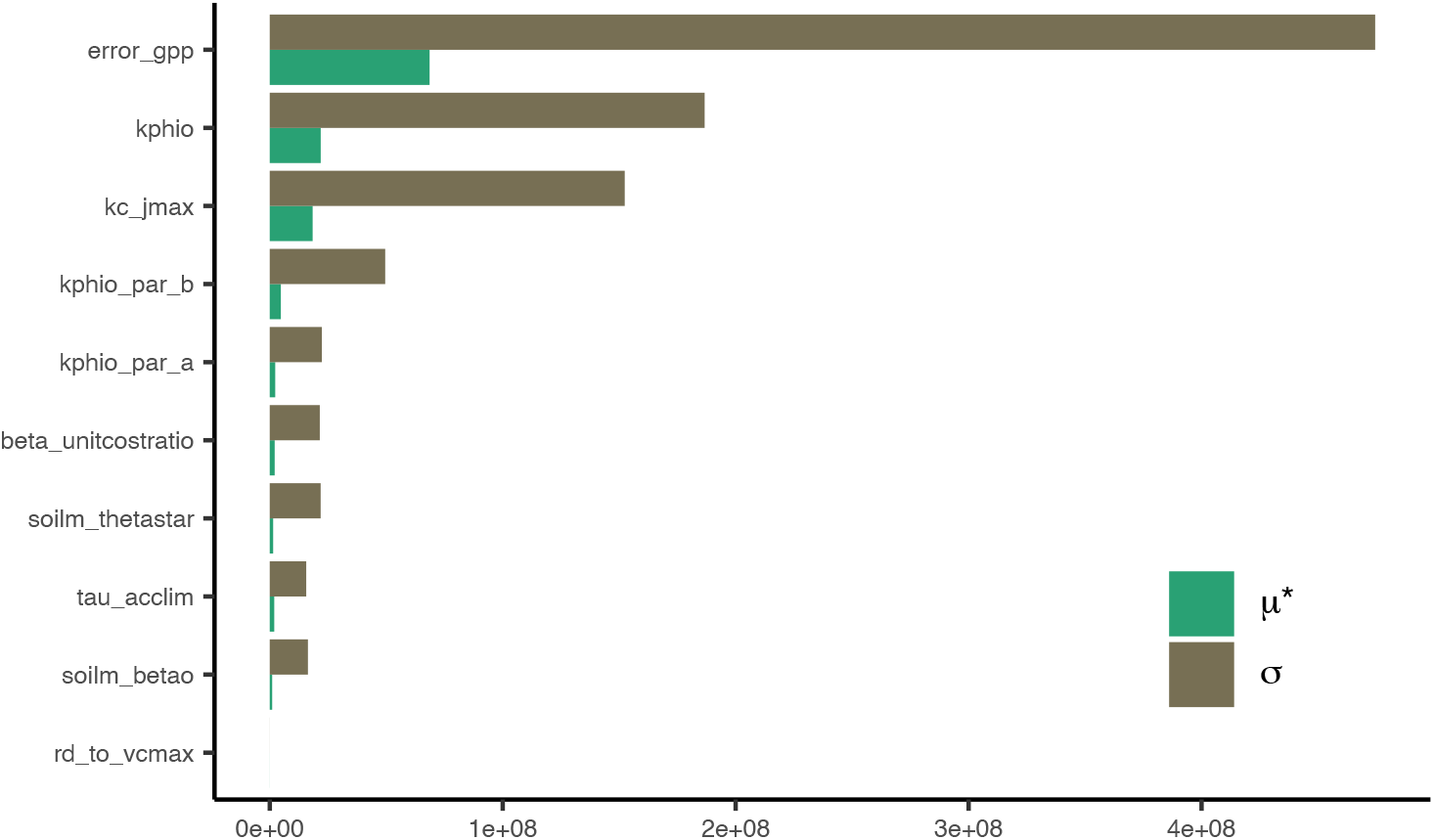
Results from the Morris sensitivity analysis. The y-axis represents all nine model parameters and the Gaussian error standard deviation (error_gpp) and the x-axis the values of the statistics *μ*\* and *σ* (unitless, like the log-likelihood). *μ*\* indicates the magnitude of the overall influence of a certain parameter on the P-model output, while *σ* measures the heterogeneity of such influence across the parameter space. The parameter names are described and identified with corresponding symbols in Tab. 2.

### 4.2 Bayesian calibration

We simultaneously calibrate a subset of the model parameters that have been identified as particularly influential (Sec. 4.1.). These include the model error term, *φ*_0_, *a, b, θ*^*^ and *β*_0_. We use the Differential-Evolution MCMC zs (DEzs) sampler (Ter Braak and Vrugt, 2008), implemented in {BayesianTools} (Hartig et al., 2023), to estimate the posterior distribution of the calibrated parameters (Fig. 3). All parameters are given a uniform prior with bounds informed by their physical interpretations (Tab. 2). The prediction error is assumed to be normally distributed, as in the Morris analysis. On a 12th Gen Intel Core i7-1270P processor, it took 1100 sec. to run 3 independent MCMC chains of 24000 iterations (of which 12000 are discarded as burn-in period). The algorithm converged with a scale reduction factor (Gelman and Rubin, 1992) of 1.05 (≤1.1). More detail, examples, and explanations of calibration diagnostics are provided through the example vignettes in the package documentation or can be inferred directly from archived the scripts used to create the results (for both, see Sections on Code availability and Data availability). The calculation of the log-likelihood is implemented in the function ‘cost_likelihood_pmodel()’, enabling custom calibrations, also for multiple target variables that are considered simultaneously during model calibration (examples also provided in the package documentation).

**Figure 3:**
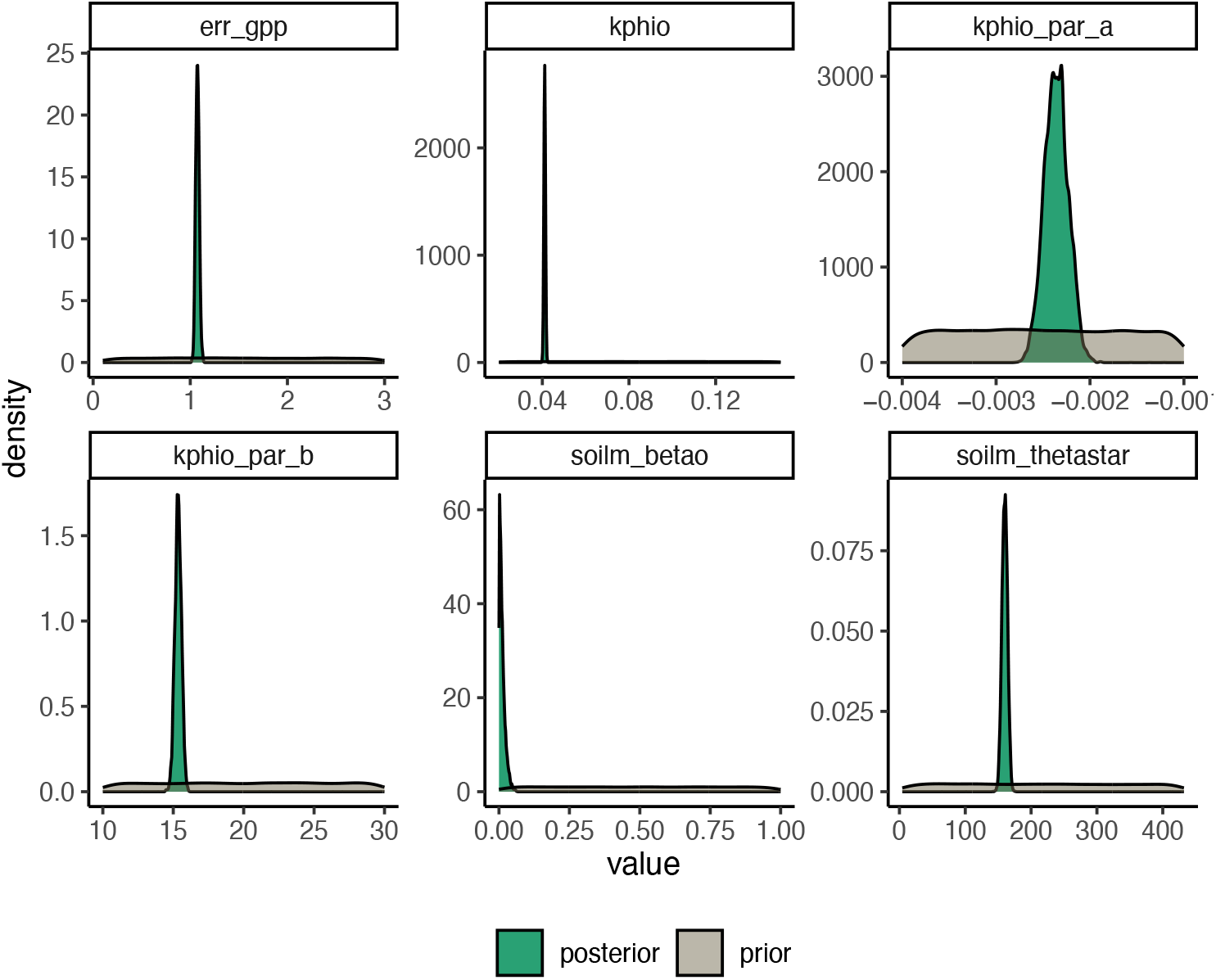
Prior and posterior distributions of the calibrated model parameters and error term. The maximum a posteriori (MAP) estimate for the gaussian error standard deviation is 1.06 and the estimate for each calibrated parameter are given in Tab. 2.

### 4.3 Inference and prediction uncertainty estimation

The parameter sets generated by the MCMC chains provide the basis for inference (model prediction) and prediction uncertainty estimation, allowing us to get insights into the sources of uncertainty. We consider a simple representation of the uncertainty, split between the parameter uncertainty and the model error (Dietze, 2017).

Retaining 600 samples from the combined Markov chains, statistically representative of the joint parameter posterior distribution (estimated during the calibration), we ran the P-model for each set of parameters to predict GPP. The credible interval was computed for each time step from the posterior distribution of predicted GPP. The prediction interval for GPP was computed by adding the Gaussian error standard deviation error, to the predicted GPP. Fig. 4 shows that the uncertainty ascribable to the parameters (in green) is much smaller than the uncertainty due to the model error (orange area). A point estimate of GPP at each time step is calculated as the median of the posterior distribution of GPP (dark orange line).

**Figure 4:**
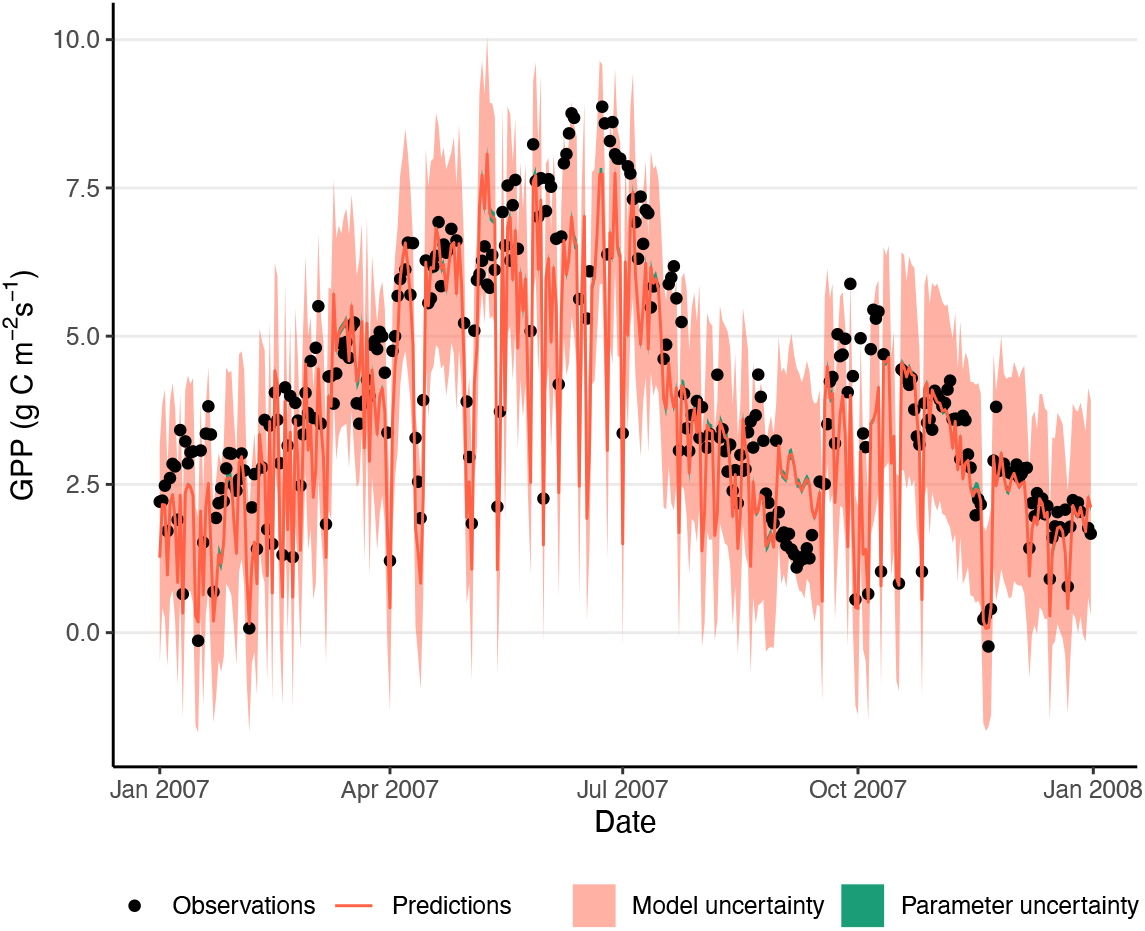
Predicted and observed daily mean GPP. The comparison is provided for the first year of GPP observations (black dots) at the site FR-Pue against GPP predictions (red line), calculated as the median of the posterior distribution. A light green band indicates the 90% credible interval for GPP predictions, which captures parameter uncertainty, while the 90% predictive interval for GPP predictions (in orange) captures model uncertainty.

## 5 Discussion

The {rsofun} R package provides a user-friendly and fast implementation of the P-model. It implements off-the-shelf model-data assimilation routines with simple function calls, while maintaining flexibility for future experiments and further development of model uncertainty estimation. The {rsofun} vegetation modelling framework is designed to strike a balance between integration and flexibility, enabled by extensible and user-defined calibration specifications and the parallelizable and fast low-level code implementation in Fortran. Its high computational efficiency offers the potential for effective model parameter and uncertainty estimation using Bayesian statistical methods - as demonstrated here.

With the function ‘calib_sofun()’, {rsofun} provides a blueprint for model-data assimilation and an implementation of the likelihood function with flexibility in selecting among a predefined list of model parameters and target observations. We have demonstrated here how ecosystem flux measurements of GPP from a single site can be used to estimate model parameters of the P-model and generate estimations of prediction uncertainty. The approach taken here (as in previous studies with the P-model) was to specify latent parameters directly, based on independent observations. In (Wang et al., 2017), *β* and *c*^*^ were determined from observations of *V*_cmax_, *J*_max_, and *c*_i_*:c*_a_. These were then used as constants for modelling GPP, while additional parameters related to empirical photosynthesis stress parameterizations were calibrated to GPP observations. Exploratory analyses (not shown) indicated that complementary observational constraints are necessary when calibrating *β* and *c*^*^ simultaneously with other model parameters. We have thus followed a simplified setup - used here for demonstration purposes. Future applications may use *V*_cmax_, *J*_max_, or *c*_i_*:c*_a_ data directly as additional calibration targets to provide such complementary constraints. Note also that the FvCB photosynthesis model contains additional parameters (see Tab. A2 in (Stocker et al., 2020)) that are treated as constants here - as in previous publications (Bloomfield et al., 2023; Mengoli et al., 2022; Stocker et al., 2020; Tan et al., 2021; Wang et al., 2017).

Our model implementation as an R package takes inspiration from the {r3PG} forest model (Trotsiuk et al., 2020), and our implementation of model-data integration on the basis of ecosystem data serves similar, yet reduced, aims and functionalities compared to PEcAn (https://pecanproject.github.io/index.html) (LeBauer et al., 2013). {rsofun} is designed to be minimally reliant on package dependencies and connections to specific data, while limiting the scope to a predefined set of process models (currently P-model, BiomeE at an experimentation stage). Note that the {rpmodel} R package (available on CRAN) also provides an implementation of the P-model but is written fully in R and in the form of a function of a given environment - without a treatment of temporal dependencies and without the functionalities for model-data integration.

## 6 Conclusions

We have demonstrated how important parameters contributing to uncertainty in GPP predictions by the P-model can be estimated using observations of ecosystem-level photosynthetic CO_2_ uptake flux time series. The Bayesian approach to model-data integration enables a probabilistic prediction of GPP and estimation of model parameters. The {rsofun} model implementation as an R package makes it possible to leverage a set of methods and complementary libraries for parameter estimation, sensitivity analysis, and calibration diagnostics - as demonstrated here. Its low-level code in Fortran is geared towards computational efficiency. Provision of {rsofun} as an open-access library aims at lowering the bar of entry to vegetation modelling for both field ecologists and computational ecologists and serves as an Open Science resource for future model development and experimentation and the further development of model uncertainty estimation.

## Appendix A: Calibrated empirical functions

A new formulation of the temperature dependency of the quantum yield efficiency *φ*_0_ is introduced in the {rsofun} package, allowing more flexibility than Eq. 18 in (Stocker et al., 2020). It is expressed as follows:

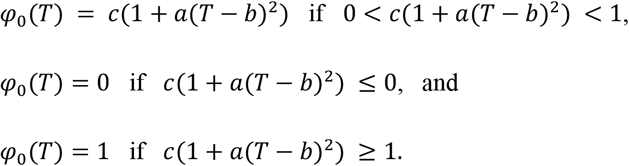

Where *T* stands for temperature, *c* is the quantum yield efficiency at optimal temperature (in mol mol ^!#^), *a* is a unitless shape parameter and *b* is the optimal temperature. Whenever a = 0, the quantum yield efficiency is kept constant at φ_$_ = c. A possible improvement for the model would be to use a peaked Arrhenius function instead of a parabola (Medlyn et al., 2002). Furthermore, the soil moisture stress function follows (Stocker et al., 2020), but the parameters considered for calibration there differ from the calibratable parameters in the package.

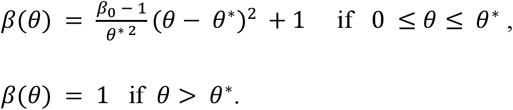

In the equation above, θ stands for the plant-available soil water (in mm) and θ^∗^ for the threshold indicating when the plants start being water stressed. The intercept β_$_ is the β reduction at low soil moisture, that is β_$_ = β(0). This intercept is now calibrated directly, rather than expressed as a function of mean aridity (see eq. 20 in (Stocker et al., 2020)).

### Code availability

The {rsofun} R package can be installed from CRAN (https://cran.r-project.org/package=rsofun) or directly from its source code on GitHub (publicly available at https://github.com/geco-bern/rsofun under an AGPLv3 licence). Versioned releases of the GitHub repository are deposited on Zenodo (https://doi.org/10.5281/zenodo.14264892). Code to reproduce the analysis and plots presented here is contained in the repository (subdirectory ‘analysis/’) and is demonstrated on the model documentation website (https://geco-bern.github.io/rsofun/, article ‘Sensitivity analysis and calibration interpretation’).

### Data availability

The model forcing and evaluation data is based on the publicly available FLUXNET2015 data for the site FR-Pue, prepared by FluxDataKit v3.4.2 (10.5281/zenodo.14808331), taken here as a subset of the originally published data for years 2007-2012. It is accessible through the {rsofun} R package and contained as part of the repository (subdirectory ‘data/’) as CSV and as files. Outputs of the analysis presented here are archived in the ‘analysis/paper_results_files/’ subfolder.

### Author contribution

B.D.S. wrote the Fortran code and designed the study. K.H. initiated the implementation of the calibration procedure and wrote the initial draft of the manuscript. J.A.P. documented the package, implemented the Bayesian calibration, sensitivity analysis and uncertainty estimation, and contributed to manuscript writing. M.M. and F.B. prepared the package for publication to CRAN and finalized the manuscript.

### Competing interests

The authors declare that they have no conflict of interest.

## Acknowledgements

We thank Florian Hartig for advice on uncertainty modelling and the maintenance of the BayesianTools package, Volodymyr Trotsiuk for an initial template of the R package, and Colin Prentice for comments on the manuscript. B.D.S. was funded by the Swiss National Science Foundation grant PCEFP2_181115. This work is a contribution to the LEMONTREE (Land Ecosystem Models based On New Theory, obseRvations and ExperimEnts) project, funded through the generosity of Eric and Wendy Schmidt by recommendation of the Schmidt Futures program (K.H., B.D.S.).

